# Novel human cell expression method reveals the role and prevalence of posttranslational modification in non-muscle tropomyosins

**DOI:** 10.1101/2021.04.05.438513

**Authors:** Peter J. Carman, Kyle R. Barrie, Roberto Dominguez

## Abstract

Biochemical studies require large protein quantities, which are typically obtained using bacterial expression. However, the folding machinery of bacteria is inadequate for many mammalian proteins, which additionally undergo posttranslational modifications (PTMs) that bacteria, yeast, or insect cells cannot perform. Many proteins also require native N- and C-termini and cannot tolerate extra tag amino acids for function. Tropomyosin, a coiled coil that decorates most actin filaments in cells, requires both native N- and C-termini and PTMs, specifically N-terminal acetylation, to polymerize along actin filaments. Here, we describe a new method that combines native protein expression in human cells with an intein-based purification tag that can be precisely removed after purification. Using this method, we expressed several non-muscle tropomyosin isoforms. Mammalian cell-expressed tropomyosins are functionally different from their *E. coli*-expressed counterparts, display multiple types of PTMs, and can form heterodimers. This method can be extended to other proteins, as demonstrated here for α-synuclein.

## Introduction

Biochemical and structural studies require large amounts of pure proteins and protein complexes. When these cannot be purified from natural sources, scientists typically opt for protein expression in bacteria, and N- or C-terminal affinity tags are commonly used for protein purification. These tags are often left on proteins during subsequent studies and, when they are enzymatically removed, extra amino acids often remain (1). While these approaches have proven powerful and will likely continue to be used in the future, there are many circumstances in which they fail to produce accurate results. For example, most human proteins undergo multiple types of posttranslational modifications (PTMs), including acetylation, phosphorylation, and methylation (2). Such PTMs play crucial roles in the regulation of protein function, including protein stability, protein-protein interactions, and enzymatic activity (3-6). Most PTMs present in human proteins cannot be processed by expression systems such as bacteria, yeast, or insect cells (7-12). These expression systems may also lack the adequate machinery to fold certain mammalian proteins (13-16). Many proteins also require native N- and C-termini and cannot tolerate the presence of extra amino acids from purification tags for proper function (17-21). We were confronted with these problems while studying tropomyosin (Tpm), a protein of central importance in the actin cytoskeleton (22).

Tpm is a coiled coil dimer that polymerizes head-to-tail to form two contiguous chains on opposite sides of the actin filament (F-actin). Through alternative splicing of four different genes (*TPM1-4*), up to 40 distinct Tpm isoforms can be produced, of which ∼30 have been detected as proteins (23). Most isoforms are ∼280-amino acids long (called long isoforms), comprising seven pseudo-repeats of ∼40 amino acids, with each pseudo-repeat spanning the length of one actin subunit along the long-pitch helix of the actin filament. Long Tpm isoforms decorate the contractile actin filaments of muscle cells. In non-muscle cells, the majority of actin filaments are also decorated with Tpm (24). Some of the non-muscle Tpm isoforms lack exon 2 and as a consequence are one pseudo-repeat shorter (called short isoforms). Because the six human actin isoforms are very similar, sharing 93–99% sequence identity, it has been proposed that the decoration of actin filaments with distinct Tpm isoforms provides a mechanism for filaments to assume different identities and become functionally segregated in cells (25). Tpm can move azimuthally on actin, helping to regulate the interactions of most actin-binding proteins, including myosin (26,27).

Although there is high sequence similarity among Tpm isoforms, gene knockout and isoform overexpression studies have shown that they play non-redundant roles (22). For example, Tpm4.2 knockout causes low platelet count (28), whereas Tpm3.5 knockout causes softer, less mechanically resilient lenses in the eye (29), indicating that other Tpm isoforms cannot compensate for the lack of Tpm4.2 or Tpm3.5. Similarly, different Tpm isoforms have been implicated in processes such as muscle contraction (30,31), cytokinesis (32,33), embryogenesis (34,35), and endocytosis (36). The temporal expression of Tpm isoforms is also tightly controlled. Thus, sixteen isoforms are up- or down-regulated at different stages of brain development in rodents (37,38). Tpm3.1 and Tpm3.2 are found in the axon in developing neurons, whereas in mature neurons they are expressed only in cell bodies and replaced in axons by Tpm1.12 and Tpm4.2 (39).

PTMs, and in particular N-terminal acetylation (Nt-acetylation) (40-42) and phosphorylation (43-46), have been shown to critically regulate Tpm function. Proper head-to-tail assembly of Tpm coiled coils along actin filaments requires native N- and C-termini, i.e. free of extra amino acids from tags and Nt-acetylated (40,47), whereas the phosphorylation state of Tpm can also modulate the stiffness and affinity of the head-to-tail interaction as well as the F-actin binding affinity (43,44,46). Despite the importance of these factors, previous biochemical studies have focused on one or two Tpm isoforms purified from muscle or *E. coli*-expressed full-length and peptide fragments of Tpm isoforms that lack PTMs and native N and C-termini. While extra amino acids have been added at the N-terminus to substitute for Nt-acetylation (40,41,48,49), it is unclear how precisely this approach mimics acetylation. Tpm coexpression with the yeast acetyltransferase NatB in *E. coli* has been also used to acetylate the N-terminus (50). However, while yeast NatB can acetylate the N-terminus, the extent of this modification varies for different Tpm isoforms, being as low as 30% for certain mammalian Tpm isoforms, whereas other types of PTMs are not accounted for (8). Furthermore, as we show here, the initiator methionine of many Tpm isoforms must be removed by mammalian N-terminal aminopeptidases, and Nt-acetylation occurs on the second residue, which this approach can also not account for (51).

Here, we describe a method that combines expression in human Expi293F cells, which grow in suspension to high density and contain the native PTM machinery, with an intein-based affinity tag that can be precisely removed through self-cleavage after purification. We applied this method to the expression of six Tpm isoforms. We demonstrate the general applicability of this method to other proteins by also showing expression of α-synuclein, a protein implicated in several neurodegenerative disorders whose function is critically regulated by numerous PTMs (52,53). We characterize the expressed proteins, including a functional analysis of the expressed Tpm isoforms in comparison with their *E. coli*-expressed counterparts, and a comprehensive analysis of PTMs using proteomics analysis. We finally show that non-muscle Tpm isoforms can form heterodimers *in vitro* and in cells, as previously shown for muscle isoforms purified from tissues.

## Results

### Human cell expression and intein-mediated purification method and its application to Tpm and α-synuclein

The expression method developed here combines mammalian cell expression with a self-cleavable affinity tag. The cells used are Expi293F, commercialized by Thermo Fisher Scientific, which are based on human embryonic kidney (HEK) cells and have been adapted to grow in suspension to high density (54,55). These cells can be transiently transfected, display high protein expression levels, and have the endogenous protein folding and PTM machineries necessary for native expression and modification of human proteins. The self-cleavable affinity tag consists of a modified intein element followed by a chitin-binding domain (CBD), and is based on the IMPACT (intein-mediated purification with an affinity chitin-binding tag) system marketed by New England Biolabs (56,57). Both, N- and C-terminal intein-CBD tags are commercially available for bacterial protein expression, but we opted here to use a C-terminal intein-CBD tag for expression in mammalian cells to allow for N-terminal processing and acetylation, which affects ∼90% of human proteins (58). After purification on the high-affinity chitin resin, self-cleavage of the intein takes place precisely after the last endogenous amino acid of the expressed target protein, leaving no extra amino acids from the tag. While both mammalian cell expression (59-61) and intein-CBD purification tags (62) have been previously used, to our knowledge this is the first time that the two are combined in a single method. Self-cleavage of the intein is thiol-catalyzed, meaning that a portion of the intein of bacterial origin is prematurely activated in the reducing environment of the cytoplasm of eukaryotic cells, likely explaining why they have not been used for mammalian cell expression.

We built a new vector (named pJCX4), which incorporates the *Mxe* GyrA intein and CBD into a mammalian-cell expression vector (*Figure 1A*). The *Mxe* GyrA intein is a 198-amino acid self-splicing protein from *Mycobacterium xenopi* (63). In its native form, the intein catalyzes its own excision from the precursor protein, forming a new peptide bond between the N- and C-terminal fragments that flank the intein (64,65). To harness this process for expression and purification of a target protein, the N- and C-terminal fragments of the intein precursor protein are substituted respectively by the target protein and the CBD affinity tag, and two amino acids at the C-terminus of the intein moiety (Asn-Cys) are mutated to Ala-Thr, which turns the intein splicing activity into precise cleavage at the junction between the target protein and the intein (66).

**Figure 1.**
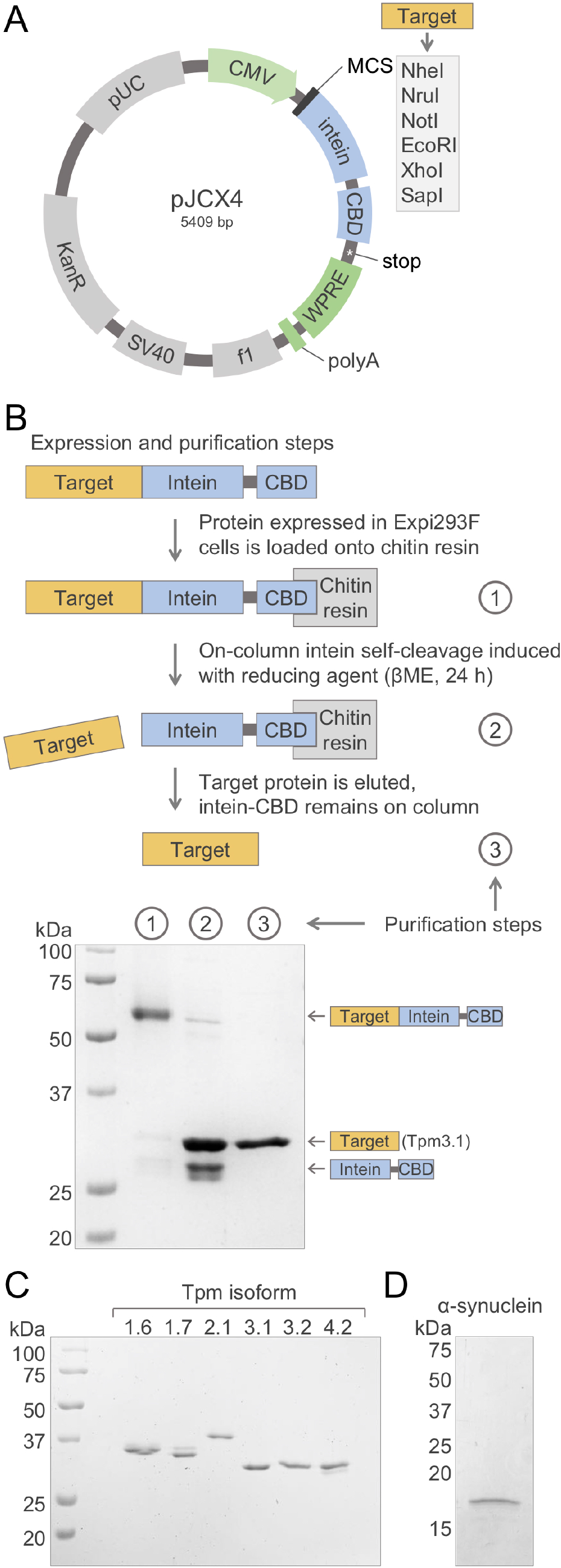
Cloning, expression and purification of Tpm isoforms and α-synuclein in Expi293F cells. (A) Schematic representation of the mammalian-cell expression vector designed here, featuring an intein-CBD affinity-purification and self-cleavage element. Cloning into this vector results in the expression of the target protein as a fusion protein with intein-CBD (see also *Figure S1*). (B) Schematic representation of the expression and purification steps, including purification on a chitin resin of the protein expressed in Expi293F cells, thiol-catalyzed self-cleavage of the intein (using β-mercaptoethanol or βME), and elution of the target protein. As an example, SDS-PAGE analysis at the bottom of the panel illustrates the progression of these steps for Tpm3.1 (see also *Figure S1*). (C and D) SDS-PAGE analysis and Coomassie Brilliant Blue R-250 (BIO-RAD) staining of the six Tpm isoforms (panel C) and α-synuclein (panel D) expressed and purified using this strategy, and further purified using ion exchange chromatography (see Materials and methods).

In addition to the intein-CBD element, the expression vector includes the cytomegalovirus promoter (CMV) and the simian virus 40 (SV40) polyadenylation (polyA) signal, as well as the woodchuck hepatitis virus posttranscriptional regulatory element (WPRE) for enhanced expression (*Figure 1A*). It also includes pUC and f1 origins of replication for propagation in *E. coli* and the SV40 origin of replication for propagation in mammalian cells. Finally, the vector contains the kanamycin resistance gene for selection in bacteria. These elements collectively allow for plasmid cloning and amplification in *E. coli* and expression in Expi293F cells. A number of restriction sites are available to clone the 5’ end of a target gene, while the SapI or SpeI sites must be used to clone the 3’ end, to ensure that no vector- or tag-derived amino acids remain after self-cleavage of the intein (*Figure 1A, Figure S1A and B*). Growth and transfection of Expi293F cells is carried out according to the manufacturer’s protocol, with minor changes (see Materials and methods). After harvesting and lysing cells, the fusion precursor protein is loaded onto a chitin affinity resin. The high affinity of the CBD-chitin interaction allows for extensive washing, and the resin can be analyzed during washing by SDS-PAGE to monitor the removal of contaminants (*Figure 1B, Figure S1C*). The intein is then activated with the addition of a reducing agent which allows the target protein to be released from the column.

**Figure S1.**
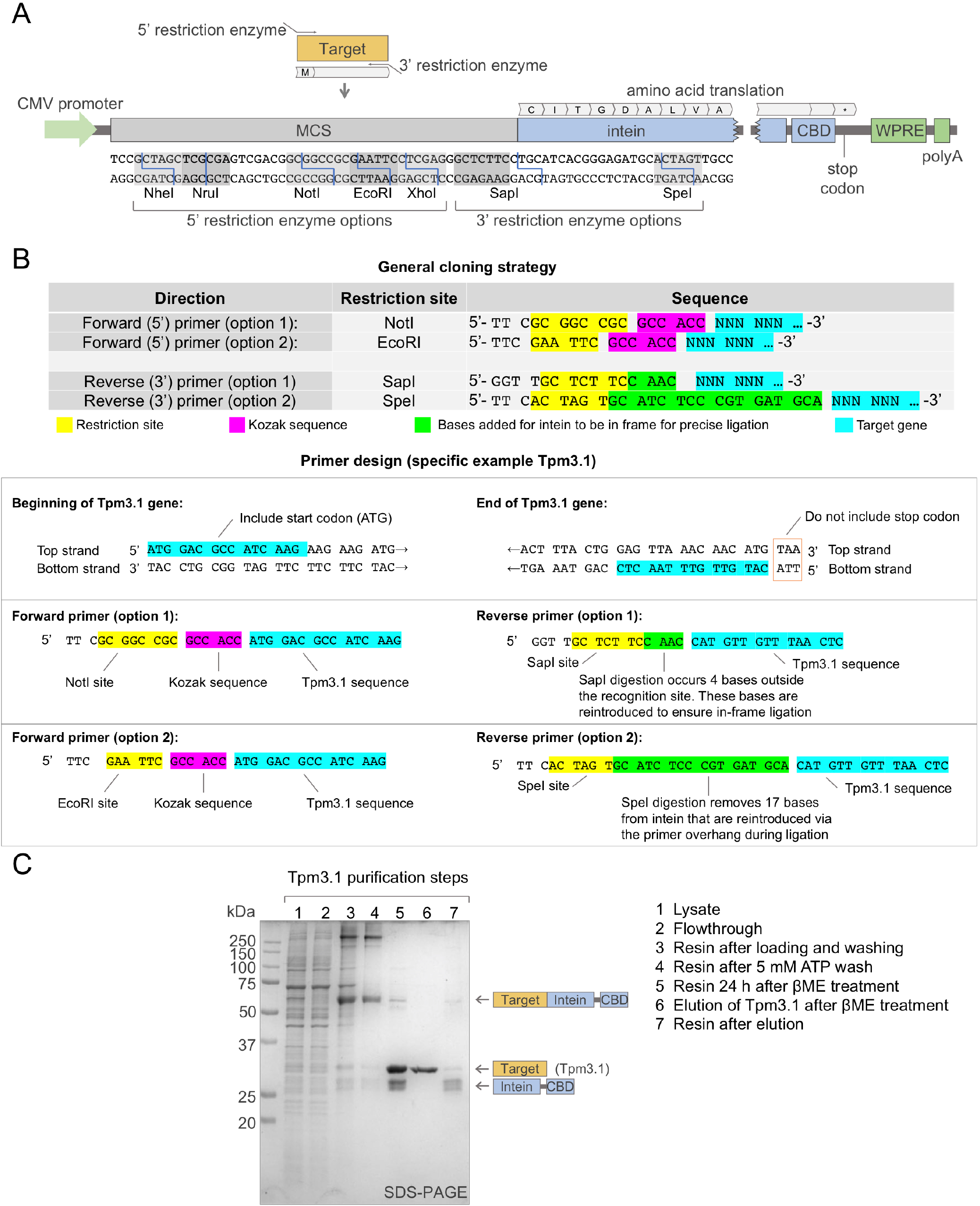
Instructions for cloning of a target gene into vector pJCX4. (A) Schematic representation of the expression cassette of vector pJCX4 for mammalian cell expression, including a restriction enzyme map of the multicloning site for the insertion of a target gene (see also Figure 1A). (B) Two examples of forward (5’) and reverse (3’) primer design to amplify a target gene for insertion into vector pJCX4, showing the general strategy (above) and a specific example for Tpm3.1 (below) (C) SDS-PAGE characterization and Coomassie Brilliant Blue R-250 (BIO-RAD) staining of the purification steps for Tpm3.1.

We used this method here to express and purify the six most studied Tpm isoforms and α-synuclein (*Figure 1C and D*). As an example, after extensive washing on the chitin resin, the fusion protein Tpm3.1-intein-CBD is nearly contaminant free (step 1 in *Figure 1B*). Self-cleavage of the intein, induced with β-mercaptoethanol for 24 h (step 2), is followed by elution of the target protein (step 3). Although the proteins were mostly pure after the intein-affinity purification step, most proteins here were additionally purified by ion exchange chromatography. Final yields varied from protein to protein, ranging from 0.25 to 2 mg of purified proteins from 250 mL of Expi293F culture.

### Post-translational modification of Expi293F-expressed Tpm and α-synuclein

To evaluate the presence of PTMs and native N- and C-termini in the proteins expressed here we used two approaches, tandem mass spectrometry (MS)-based proteomics and Pro-Q Diamond phosphoprotein gel staining (Thermo Fisher Scientific). Fluorescent Pro-Q Diamond staining is used to directly detect the presence of phosphate groups on tyrosine, serine, or threonine residues in acrylamide gels. It is important to note that signaling cascades that get activated during cell lysis can lead to increased dephosphorylation (67-69). Therefore, phosphatase inhibitors can be used during cell lysis if necessary to preserve native phosphorylation sites (70,71). We used this approach here in the Pro-Q Diamond staining analysis (*Figure 2B and D*), but not for PTM proteomics analysis.

**Figure 2.**
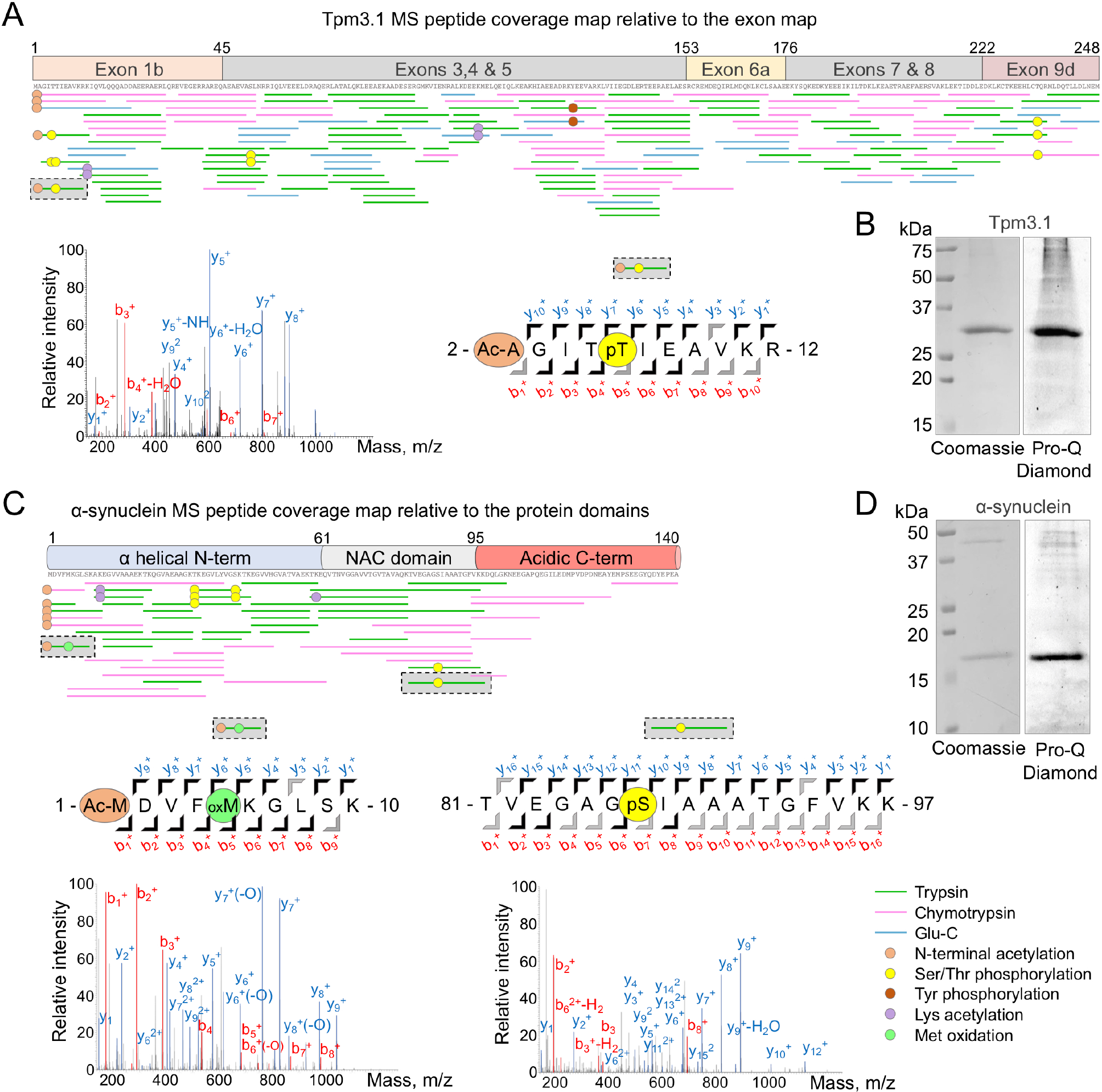
Post-translational modification analysis of Expi293F-expressed Tpm3.1 and α-synuclein. (A and C) Mass spectrometry peptide coverage maps of Tpm3.1 and α-synuclein, shown relative to exon and protein domain diagrams, respectively. Green, magenta and blue lines indicate peptides obtained by digestions with trypsin, chymotrypsin, and Glu-C, respectively, and whose masses were identified by mass spectrometry using a false discovery rate (FDR) threshold of < 1% (see also *Table S1* and *Figure S2*). Orange, yellow, red, purple, and green circles indicate Nt-acetylation, Ser/Thr phosphorylation, Tyr phosphorylation, Lys acetylation, and Met oxidation, respectively. The mass spectrometry spectra (MS2) of three representative peptides (highlighted by black dashed lines and gray background) are shown as examples, including N-terminal peptides of Tpm3.1 and α-synuclein, and a central peptide of α-synuclein. These peptides show different types of PTMs (Nt-acetylation, Ser/Thr phosphorylation, and Met oxidation). The MS2 spectra show the b (N-terminal) and y (C-terminal) fragment ions detected by mass spectrometry for each peptide. Sequence diagrams corresponding to the three spectra illustrate all the theoretical fragment ions colored black (observed experimentally) or gray (non-observed). (B and D) SDS-PAGE analysis of purified samples of Tpm3.1 and α-synuclein, using phosphatase inhibitors to reduce sample dephosphorylation during purification, and stained using either Coomassie Brilliant Blue R-250 (BIO-RAD) or Pro-Q Diamond Phosphoprotein Gel Stain (Thermo Fisher Scientific).

To ensure full peptide coverage during MS analysis, the expressed proteins were subjected to digestions using three different enzymes: trypsin, chymotrypsin, and Glu-C. To establish the identity of each peptide in the observed MS spectra, their masses were matched to the peptides in a Tpm sequence database, using a false discovery rate (FDR) threshold of less than 1% (*Table S1*). Full peptide coverage was obtained for all the proteins expressed, including Tpm isoforms and α-synuclein, with one exception, Tpm4.2, for which coverage was 99% (*Figure 2A and C, Figure S2*). This analysis revealed several interesting findings. All the proteins, including α-synuclein, were Nt-acetylated, which as mentioned above is a precondition for optimal Tpm polymerization along actin filaments (40,41). The three short Tpm isoforms analyzed (Tpm3.1, Tpm3.2, Tpm4.2) were acetylated on the second residue. In other words, for these three isoforms the initiator methionine is first removed during N-terminal processing and Nt-acetylation then takes place on the second residue. Different N-terminal acetyl transferases (NATs) likely acetylate Tpm isoforms at the initiator methionine and at position two. Based on the known sequence specificity of the various human NATs (72), we anticipate isoforms Tpm3.1, Tpm3.2, and Tpm4.2 with N-terminal sequence Met-Ala-Gly have their initiator methionine removed by MetAP (methionine aminopeptidase) and then become acetylated by NatA on Ala-2, whereas isoforms Tpm1.6, Tpm1.7, and Tpm2.1 with N-terminal sequence Met-Asp-Ala are acetylated by NatB on the initiator methionine. α-Synuclein, with N-terminal sequence Met-Asp-Val is also acetylated on the initiator methionine by NatB (73) (*Figure 2C*). For all the proteins expressed here, intact and native C-termini peptides were identified (*Figure 2A and C, Figure S2*), confirming that self-cleavage of the intein proceeded as expected, without the removal of any endogenous amino acids of the target protein or the presence of any extra amino acids from the tag. Numerous phosphorylation sites, including serine/threonine and tyrosine phosphorylation, were identified in the expressed proteins with an FDR < 1% (*Table 1, Figure 2, Figure S2*). The identity of these phosphorylation sites is mostly consistent with previous phosphoproteomics studies curated by PhosphoSitePlus (74). Other forms of PTMs were also identified using the stringent FDR cutoff of < 1%, including most notably several sites of lysine acetylation (*Table 1, Figure 2, Figure S2, Table S1*). Methionine oxidation was also observed, including for instance α-synuclein Met-5 (*Figure 2C, Table S1*), however this is not a native modification but rather one that occurs during sample preparation for mass spectrometry (75). Table S1 contains a complete list of all the peptides and PTMs observed.

**Figure S2.**
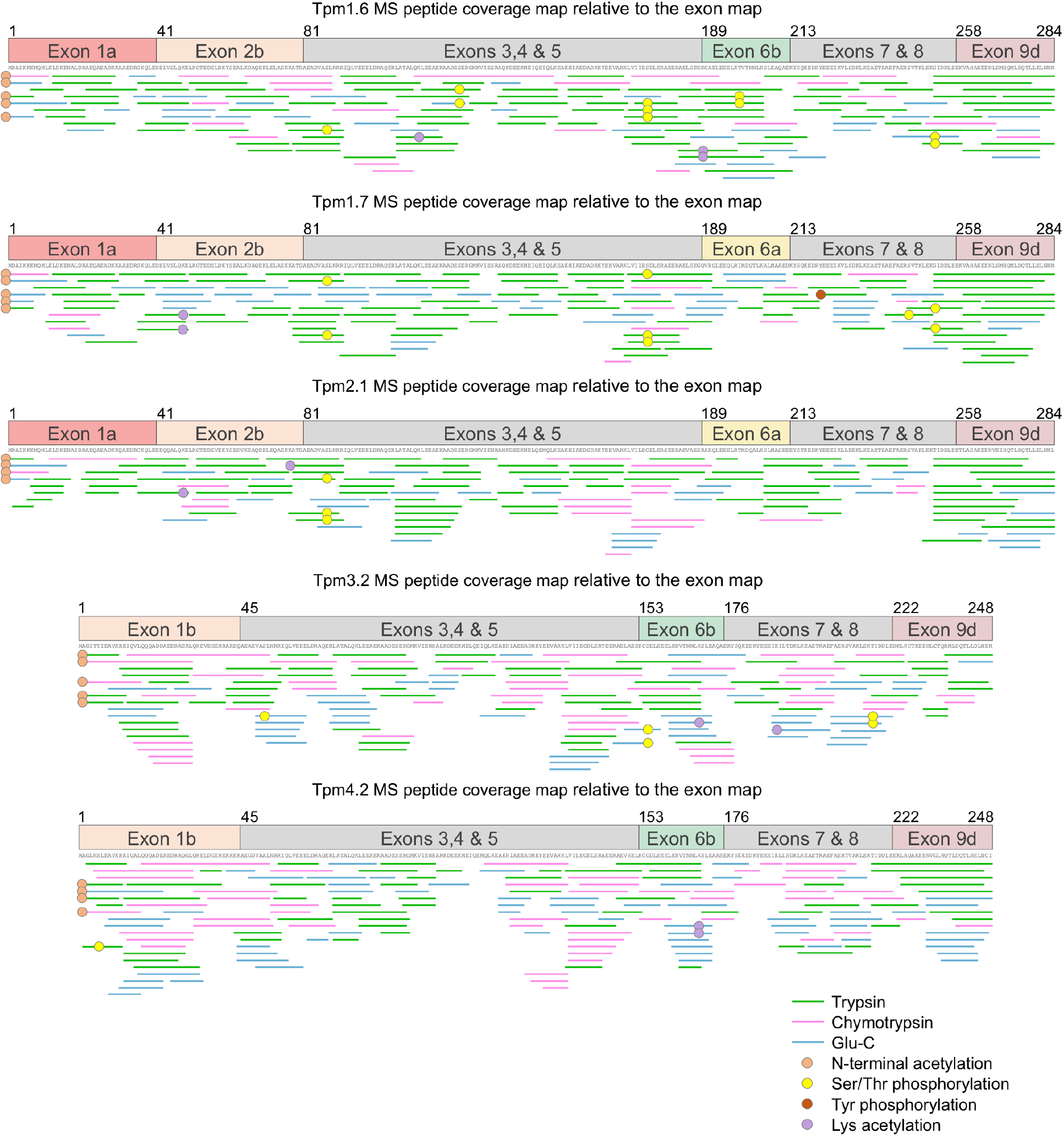
Post-translational modification analysis of Expi293F-expressed Tpm isoforms. Mass spectrometry peptide coverage maps of Tpm isoforms 1.6, 1.7, 2.1, 3.2, and 4.2, shown relative to the exon diagram of each isoform. Green, magenta and blue lines indicate peptides obtained by digestions with trypsin, chymotrypsin, and Glu-C, respectively, and whose masses were identified by mass spectrometry using a false discovery rate (FDR) threshold of less than 1% (see also Figure 2, Table S1, and Materials and methods). Orange, yellow, red, and purple circles indicate Nt-acetylation, Ser/Thr phosphorylation, Tyr phosphorylation, and Lys acetylation, respectively.

**Table 1.** PTMs identified in the human cell-expressed proteins (FDR < 1%)

| Table 1. PTMs identified in the human cell-expressed proteins (FDR < 1%) |  |  |  |  |
| --- | --- | --- | --- | --- |
| Protein | Nt-acetylation | Ser/Thr phosphorylation | Tyr phosphorylation | Lys acetylation |
| Tpm1.6 | Met-1 | Ser-87, Ser-123, Ser-174, Thr-199, Ser-252 |  | Lys-112, Lys-189 |
| Tpm1.7 | Met-1 | Ser-87, Ser-174, Ser-245, Ser-252 | Tyr-221 | Lys-48 |
| Tpm2.1 | Met-1 | Ser-87 |  | Lys-48, Lys-77 |
| Tpm3.1 | Ala-2 | Thr-5, Thr-6, Ser-51, Thr-234, | Tyr-126 | Lys-13, Lys-104 |
| Tpm3.2 | Ala-2 | Ser-51, Ser-155, Thr-216 |  | Lys-169, Lys-190 |
| Tpm4.2 | Ala-2 | Ser-6 |  | Lys-169 |
| α-synuclein | Met-1 | Thr-33, Ser-42, Ser-87 | | Lys-12, Lys-60 |

### Comparison of the activities of Expi293F-vs. *E. coli*-expressed Tpm isoforms

The defining biochemical activity of Tpm is to decorate actin filaments, which by combinatorial association of different isoforms of both actin and Tpm can give rise to a large number of Tpm-actin copolymers with distinct properties and activities in cells (25). Accordingly, we used here actin cosedimentation assays to compare the actin-binding affinities of a representative group of three Expi293F-expressed Tpm isoforms, corresponding to three different genes and comprising both short (Tpm3.1) and long (Tpm1.6 and Tpm2.1) isoforms with differently processed N-termini (see Materials and methods). Quantification of multiple such experiments, using SDS-PAGE and densitometric analysis, and fitting of the data to a Hill equation revealed the parameters of the interactions of Tpm isoforms with F-actin, including the apparent dissociation constant (*K*_*app*_) and Hill coefficient (a measure of the cooperativity of the interaction) (*Figure 3A, B and E, Figure S3*). This analysis confirmed binding of all three of the Tpm isoforms to F-actin, but with significantly higher affinity for Tpm1.6 (*K*_*app*_ = 0.32 ± 0.06 µM) than Tpm2.1 or Tpm3.1 (*K*_*app*_ = 1.78 ± 0.10 µM and 1.72 ± 0.09 µM, respectively). Interestingly, the latter two isoforms had very similar affinities for F-actin, despite one being one pseudo-repeat longer than the other and having differently processed N-termini.

**Figure 3.**
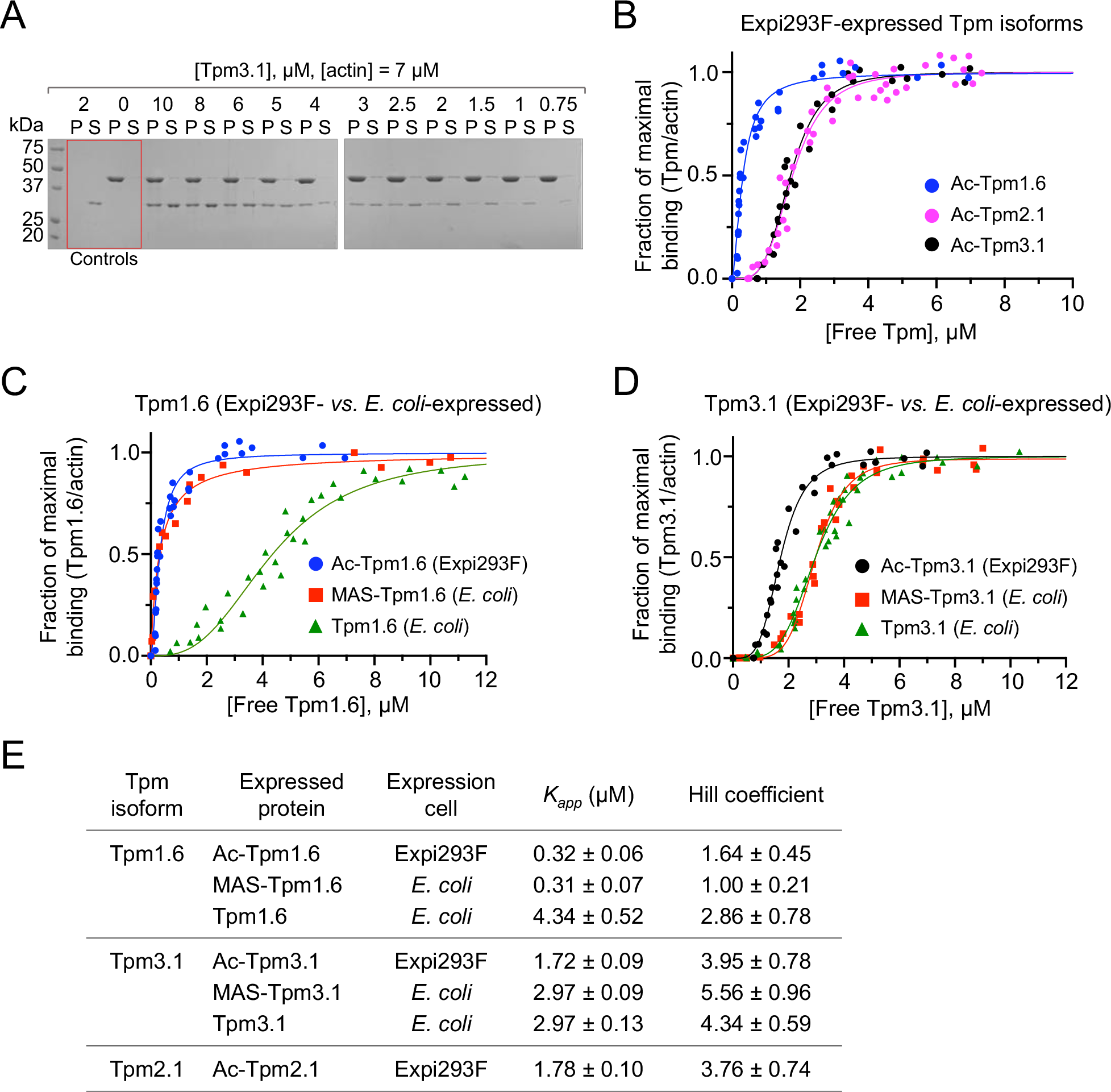
Binding of Tpm isoforms to F-actin using cosedimentation analysis. (A) Representative SDS-PAGE cosedimentation analysis of Expi293F-expressed Tpm3.1 (0.75-10 µM) with F-actin (7 µM). Controls include actin alone, found mostly in the pellet fraction (P), and Tpm3.1 alone, found exclusively in the supernatant fraction (S). (B) Between three and four cosedimentation experiments were performed for each of the three Tpm isoforms analyzed (Tpm1.6, Tpm2.1, Tpm3.1; color-coded), the density of the bands in the gels were quantified and data were plotted as the fraction of maximal binding *vs*. the free concentration of each Tpm isoform (see *Figure S3* and Materials and methods). The data were fit to a Hill equation to obtain the dissociation constant (*K*_*app*_) and Hill coefficient of each interaction. (C and D) experiments similar to those shown in panels *A* and *B* were performed to compare the F-actin-binding affinities of Expi293F-expressed Tpm1.6 and Tpm3.1 with their *E. coli-*expressed counterparts, with and without the N-terminal addition of Met-Ala-Ser (MAS), a modification thought to mimic acetylation {Gateva, 2017 #12568;Palm, 2003 #12954}. (E) Parameters of the fits shown in panels B to D, including *K*_*app*_ and Hill coefficient.

We then compared the two most studied non-muscle isoforms (Tpm1.6 and Tpm3.1) with their *E. coli-*expressed counterparts, both with and without the addition at the N-terminus of three amino acids (Met-Ala-Ser) that have been widely used to mimic acetylation (41,76). This analysis revealed at least three interesting results. First, for the long isoform Tpm1.6, the Expi293F- and *E. coli*-expressed proteins had very similar affinities for F-actin when Met-Ala-Ser was added at the N-terminus (*Figure 3C and E*). However, the affinity of *E. coli*-expressed Tpm1.6 dropped ∼15-fold when Met-Ala-Ser was not added. This result suggests that for Tpm1.6 the addition of Met-Ala-Ser adequately mimics N-terminal acetylation for binding to F-actin, although it remains to be shown whether this modification can fully account for other regulatory activities of Tpm1.6 in controlling the interactions of actin-binding proteins (ABPs) with the actin filament. Second, for the short isoform Tpm3.1 the two *E. coli*-expressed proteins, with and without Met-Ala-Ser at the N-terminus, had nearly identical affinities for F-actin, but lower than the affinity of Expi293F-expressed Tpm3.1 (*Figure 3D and E*). Therefore, for Tpm3.1 the addition of extra amino acids at the N-terminus cannot substitute for native Nt-acetylation, indicating that native PTMs added by Expi293F cells are necessary for function. We suspect this behavior may affect other Tpm isoforms, which is an important consideration since many studies have used extra N-terminal amino acids to substitute for Nt-acetylation irrespective of isoform. Third, it is known that most Tpm isoforms bind cooperatively to F-actin (77-79). Here, the isoforms that bound F-actin with lower affinity displayed higher cooperativity, and this was true among different isoforms as well as among variants of the same isoform expressed in different cell types. Thus, Tpm2.1 and Tpm3.1 exhibited higher cooperativity than Tpm1.6, whereas the *E. coli*-expressed Tpm isoforms that bound F-actin with lower affinity also showed higher cooperativity (*Figure 3E*).

**Figure S3.**
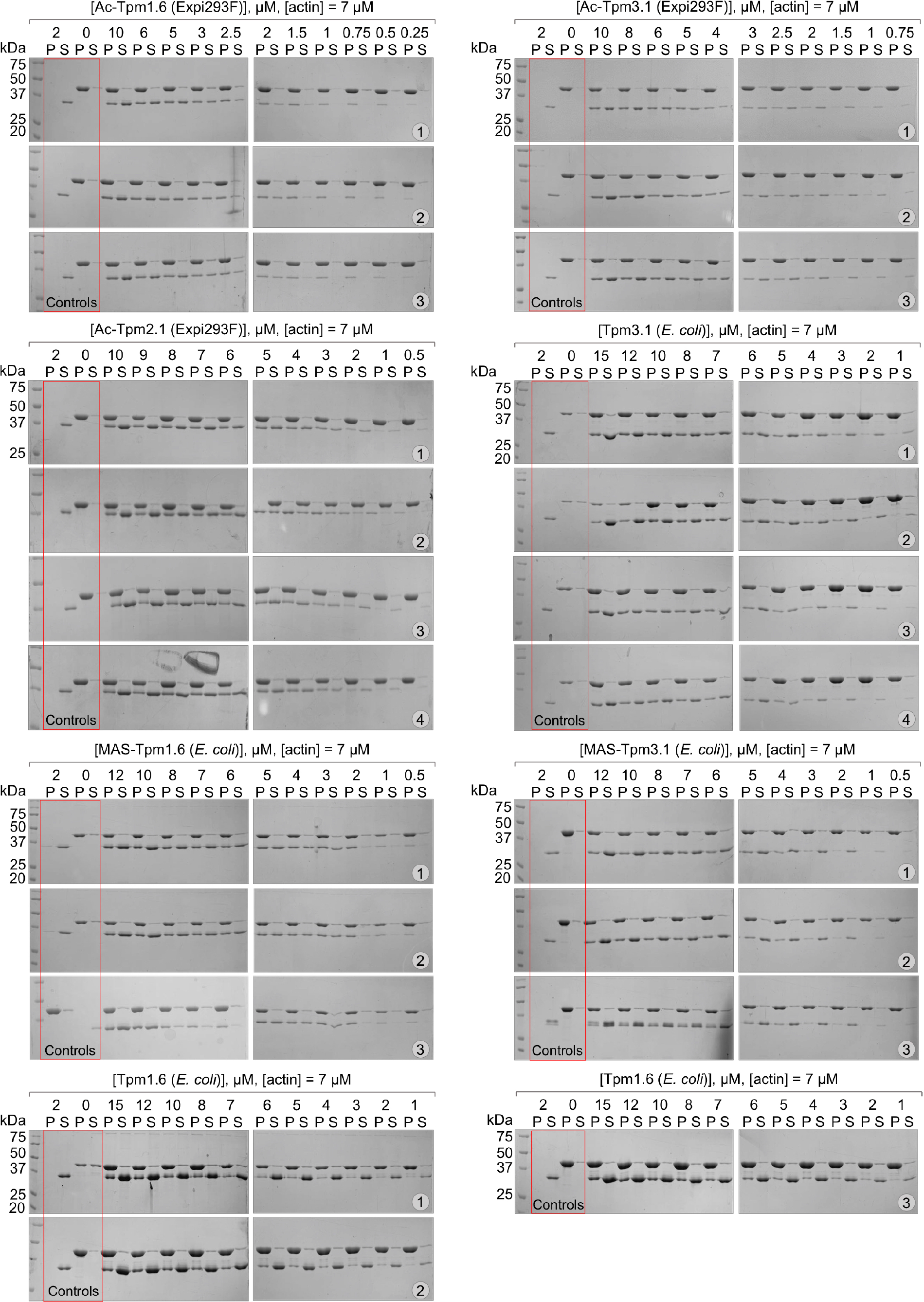
Cosedimentation of Tpm isoforms with F-actin. Cosedimentation analysis of the interactions of Tpm isoforms with F-actin using 12% SDS-PAGE gels, showing Expi293F-expressed Tpm1.6, Tpm2.1, and Tpm3.1 and *E. coli*-expressed Tpm1.6 and Tpm3.1 with and without the addition of Met-Ala-Ser (MAS) at the N-terminus to mimic Nt-acetylation. The actin concentration in all the experiments was 7 µM, whereas the concentration of Tpm varied for each experiment and is shown on the top of the gels. Controls include actin alone, found mostly in the pellet fraction (P), and Tpm alone, found exclusively in the supernatant fraction (S) for all the proteins analyzed. This data was used to produce the plots and data fits shown in Figure 3.

### Non-muscle Tpm isoforms can form heterodimers

The actin filaments of the contractile apparatus of muscle cells are decorated with two or more Tpm isoforms, which often associate as heterodimers (80-83). For instance, in skeletal muscle Tpm1.1/Tpm2.2 (also known as isoforms α and β) heterodimers form more favorably than Tpm1.1/Tpm1.1 homodimers (81). It is unknown whether non-muscle Tpm isoforms can also form heterodimers in cells.

To explore this question, we modified the mammalian cell expression system described here to coexpress pairs of Tpm isoforms. We designed a new mammalian cell expression vector, containing an internal ribosomal entry site (IRES) inserted in between two multi-cloning sites (MCS), one for each Tpm isoform being coexpressed (*Figure 4A*). The use of the coexpression IRES vector, instead of cotransfection with two separate vectors, ensures that similar amounts of the two Tpm isoforms are expressed together in time and space. Adjacent to MCS1 and MCS2 the vector contains a V5 and a FLAG affinity tag, respectively. In this way, the coexpressed Tpm isoforms have native N-termini, but either V5 or FLAG affinity tags added at the C-terminus. These tags are used for tandem affinity purification and detection via Western blot analysis (*Figure 4B*), a method we previously used to study wild-type/mutant heterodimers of IRSp53 (84). The presence of these tags likely disrupts the head-to-tail interaction of Tpm coiled coils along F-actin, but because coiled coil formation does not depend on Tpm’s ability to bind F-actin, these tags were not removed for the experiments described here. When two Tpm isoforms are coexpressed using this strategy, they can form either two independent homodimers, heterodimers or a combination of homodimers and heterodimers. To discriminate among these possibilities, we purify the mixture on a FLAG resin, wash extensively, and the eluted sample is then analyzed by anti-FLAG and anti-V5 Western blotting (*Figure 4B*). The presence of an anti-V5 signal in the eluted sample (but not in the last wash) is a strong indication of heterodimer formation. This procedure is illustrated here for the coexpressed isoform pair Tpm3.1-V5/Tpm4.2-FLAG (*Figure 4C*). Note that the anti-FLAG and anti-V5 signals disappear almost entirely in the last wash, but both signals show up strongly in the eluted sample, indicating that any Tpm3.1-V5 that was retained in the FLAG resin was forming heterodimers with Tpm4.2-FLAG.

**Figure 4.**
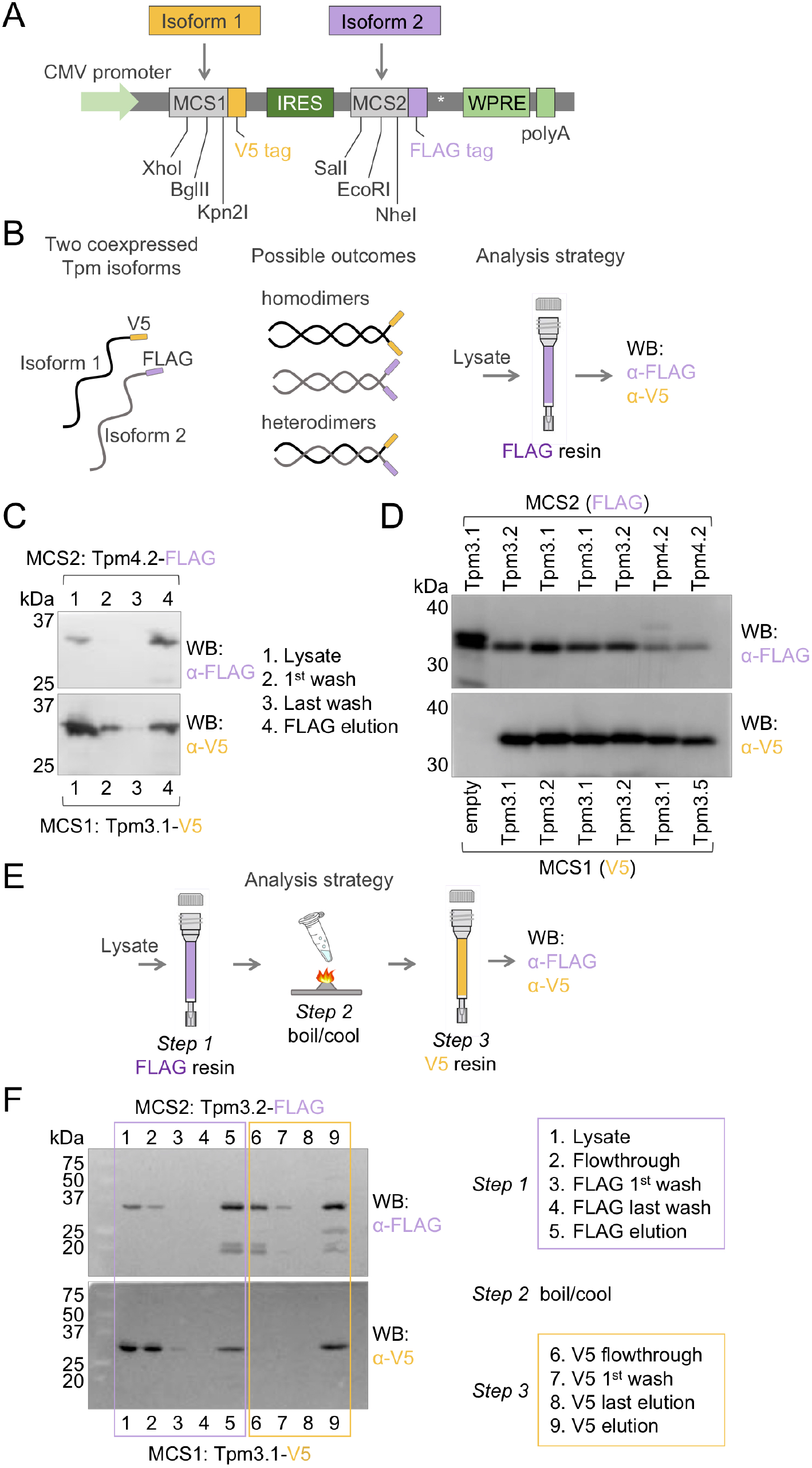
Heterodimerization of non-muscle Tpm isoforms. (A) Schematic representation of the MCS1-IRES-MCS2 elements of vector pJC8 used for coexpressio pairs of Tpm isoforms, including a depiction of the restriction sites and affinity tags (FLAG and V5) associated with each MCS. (B) Illustration of the potential asso of coexpressed Tpm pairs as homodimers or heterodimers and strategy for the FLAG tag-based purification and Western blot-based detection of such species usi anti-FLAG and anti-V5 antibodies. (C) Illustration of this strategy for the coexpressed pair of Tpm3.1-V5 and Tpm4.2-FLAG, showing the presence of both isoform heterodimers reactive to both anti-FLAG and anti-V5 antibodies, in the elution sample after extensive washing on the FLAG resin. (D) Western blot analysis of the elution samples from seven such experiments, including six Tpm pairs and a control experiment with Tpm3.1-FLAG cloned into MCS2 and no Tpm isoform cloned MCS1. (E) Experimental design for the detection of Tpm heterodimers after unfolding and refolding. Following the FLAG elution, samples are boiled and cooled a hen purified on a V5 resin. (F) Illustration of this strategy for the coexpressed pair of Tpm3.1-V5 and Tpm3.2-FLAG. After extensive washing, the Western blot de of anti-FLAG and anti-V5 signals in the sample from the V5 resin demonstrates that Tpm3.1/Tpm3.2 heterodimers formed spontaneously during refolding.

Because heterodimerization of long muscle isoforms has been demonstrated (80,81,85), the analysis here focused on four short, non-muscle Tpm isoforms: Tpm3.1, Tpm3.2, Tpm3.5, and Tpm4.2. Note that Tpm isoforms of different lengths are unlikely to heterodimerize since this would lead to parts of the hydrophobic core of the coiled coil being exposed to the solvent. We co-expressed the following pairs: Tpm3.1-V5/Tpm3.2-FLAG, Tpm3.1-V5/Tpm4.2-FLAG, and Tpm3.5-V5/Tpm4.2-FLAG, which all formed heterodimers (*Figure 4D*). To control for potential differences in the expression levels of the proteins cloned into MCS1 and MCS2, we switched the order of isoforms and coexpressed the pair Tpm3.2-V5/Tpm3.1-FLAG, which also formed a heterodimer. Another control showed that V5- and FLAG-tagged isoforms Tpm3.1 and Tpm3.2 form homodimers despite the different tags. Finally, Tpm3.1-FLAG cloned alone into MCS2 (MCS1 empty) also eluted as a homodimer and no anti-V5 signal was detected in the eluted sample (*Figure 4D*).

These results strongly suggest that Tpm3.1, Tpm3.2, Tpm3.5, and Tpm4.2, and likely other non-muscle isoforms can form heterodimers as well as homodimers in cells when coexpressed together. We then asked whether they would preferentially reassociate as homodimers or heterodimers after boiling *in vitro*, which unfolds the coiled coil, as shown for muscle isoforms (86-88). For this, we modified the procedure described above introducing two additional steps; after purification on the FLAG resin, the eluted samples were boiled, then allowed to slowly refold during cooling, repurified on a V5 resin, and then analyzed by anti-FLAG and anti-V5 Western blotting (*Figure 4E*). We analyzed in this manner the pair Tpm3.1-V5/Tpm3.2-FLAG and monitored each step of this procedure by Western blotting (*Figure 4F*). The initial steps of the procedure (i.e. purification on the FLAG resin) confirmed the formation of heterodimers, as described above (*Figure 4D*). After boiling/cooling the sample was loaded onto the V5 resin. The flowthrough from the V5 resin reacted only with the anti-FLAG antibody, indicating that some of the heterodimers had reassociated as homodimers. Yet, the V5 elution showed strong anti-FLAG and anti-V5 signals. Any Tpm3.2-FLAG retained in the V5 resin was likely forming heterodimers with Tpm3.1-V5. Together, the data presented here show that non-muscle Tpm isoforms can form both homodimer and heterodimers in cells and *in vitro*.

## Discussion

### Main features and general applicability of the expression-purification method

We have developed a novel method for protein expression and purification that integrates the advantages of mammalian cell expression with those of an intein-CBD purification tag. The mammalian cells used, Expi293F, grow to high density in suspension and are adapted for high-yield protein expression. More importantly, these cells naturally contain the necessary folding and PTM machineries to produce fully native human proteins. The intein-CBD tag, on the other hand, provides three key advantages. First, it is a high-affinity tag that allows for extensive on-column washing, such that high protein purity can be achieved in a single purification step. Second, this is a self-cleavable tag, which can be activated on-column, bypassing the necessity for enzymatic cleavage after purification and additional purification associated with other tags that can in some cases produce non-specific cleavage and introduces two new contaminants, i.e. the cleavage enzyme and the released affinity tag. Third, through proper design of the cloning primers (as shown in *Figure S1*) self-cleavage of the intein can take place precisely after the last C-terminal amino acid of the expressed protein, without the deletion of endogenous amino acids or the presence of extra amino acids from the tag. One potential drawback of this system is that a certain amount of premature cleavage may take place in the reducing environment of the cytoplasm of Expi293F cells, which ultimately reduces protein yields. Yet, for all the proteins expressed here yields of ∼1 mg of highly pure proteins were obtained from relatively small Expi293F cultures of 250 mL. Such protein amounts are clearly sufficient for most biochemical and structural studies and the technology is cost permissive for most laboratories possessing a mammalian cell incubator. The combined benefits and relative simplicity of this method make it easily adaptable for the expression of most human proteins, and particularly those whose activities depend upon key PTMs. Thus, while the initial motivation for this work was the expression of several cytoskeletal proteins of interest to our laboratory, such as Tpm, we extended it here to the expression of α-synuclein, a protein of intensive interest and a notable target of many PTMs (52,53). To facilitate the adoption of this system, we are depositing the two vectors developed here with the Addgene plasmid repository (www.addgene.org).

### PTMs of human cell-expressed non-muscle Tpm isoforms

The use of the human cell-expression system described here allowed us to make several discoveries. One of these discoveries concerns the presence of numerous forms of PTMs, including Nt-acetylation, Ser/Thr and Tyr phosphorylation and Lys acetylation in non-muscle Tpm isoforms (*Figure 2A, Figure S2, Table 1*). We have limited the list of PTMs reported here to sites detected with very high fidelity (FDR < 1%), but many other sites were detected with relatively high confidence (FDR < 5%) and could possibly play functional roles (*Table S1*). The importance of Ser/Thr phosphorylation in muscle Tpm isoforms has been recognized since the seventies (89,90), and a more recent study suggests that phosphorylation of Ser-283 near the C-terminus of striated muscle Tpm (Tpm1.1) enhances the stiffness of the head-to-tail interaction (43). Virtually nothing, however, is known about the role of Ser/Thr phosphorylation in non-muscle Tpm isoforms. Curiously, among the phosphorylation sites observed here, several occur near the N- and/or C-terminal ends of Tpm3.1, Tpm1.6, Tpm1.7, Tpm3.2, and Tpm4.2, and could possibly have a similar role as in muscle Tpm1.1, by modulating the stiffness of the head-to-tail interaction. To our knowledge, the role Tyr phosphorylation in Tpm isoforms remains unexplored, and our finding here of Tyr phosphorylation sites in Tpm3.1 and Tpm1.7 opens an interesting new line of inquire to understand the role of this modification. Lys acetylation of Tpm isoforms is also virtually unexplored. One of the Lys acetylation sites detected here occurs near the N-terminus of Tpm3.1 and could also modulate the polymerizability of this isoform along actin filaments, as *in vitro* studies have suggested for muscle Tpm (91). A recent study also finds that Lys acetylation of F-actin decreases Tpm-dependent inhibition of actomyosin activity (92). Lys acetylation of non-muscle Tpm isoforms could have a similar effect, by modulating the interactions of ABPs. The presence and extent of Nt-acetylation among non-muscle Tpm isoforms is to this day questioned (93). Since ∼90% of human proteins are Nt-acetylated (58) and Nt-acetylation plays a crucial role in the polymerization of muscle Tpm isoforms (41), it is not surprising that we found here that all the non-muscle isoforms analyzed were Nt-acetylated. What is more, we did not detect a single N-terminal peptide that was not acetylated, suggesting that N-terminal acetylation is a constitutive modification for all Tpm isoforms. More surprising, but consistent with the known sequence specificity of human NATs (72), was the observation that the short Tpm isoforms (Tpm3.1, Tpm3.2, Tpm4.2) undergo Nt-acetylation on the second residue, after removal of the initiator Met by MetAP, which the PTM machinery of Expi293F cells was also able to achieve for these overexpressed proteins. These three isoforms are likely Nt-acetylated by NatA, whereas the long isoforms (Tpm1.6, Tpm1.7, Tpm2.1) are likely Nt-acetylated by NatB on the initiator Met (72).

### Different activities of Tpm isoforms expressed in human and *E. coli* cells

The vast majority of *in vitro* studies on Tpm isoforms have used *E. coli-*expressed proteins. For years, these studies have used the addition of extra amino acids at the N-terminus of Tpm as a way to mimic Nt-acetylation (40,41,48,49) and have assumed that this modification fully replicates the natural capacity of Tpm to polymerize along actin filaments. By comparing for the first time side-by-side the F-actin-binding activity of naturally acetylated and *E. coli*-expressed Tpm isoforms ± Nt-acetylation-mimics we have shown here that this assumption is not always true. Thus, while the N-terminal addition of Met-Ala-Ser mostly restores the F-actin-binding affinity of *E. coli*-expressed Tpm1.6, it fails to do the same for Tpm3.1 (*Figure 3*). Tpm3.1 is a short isoform, whereas Tpm1.6 is a long isoform and interacts with one more actin protomer along F-actin. While this difference could possibly explain their different behaviors in these experiments, it appears prudent to test this effect individually for each Tpm isoform, particularly that this work provides a solution to the problem of Nt-acetylation. Indeed, Tpm2.1 is also a long isoform, but we found its affinity for F-actin to be very similar to that of Tpm3.1 (*Figure 3A*). We also noticed an interesting correlation between affinity and cooperativity; lower Tpm binding affinity for F-actin correlates with higher cooperativity of the interaction, both among naturally occurring isoforms or within a single isoform with different head-to-tail interaction sequences (*Figure 3E*). Although we could not find a previous reference to this correlation, it is intuitively clear that as the affinity of single Tpm dimers for F-actin becomes lower, binding to F-actin would become increasingly dependent on the pre-polymerization of Tpm into mini-filaments of ever-growing length through head-to-tail interaction. This is also why a strong head-to-tail interaction, which Nt-acetylation favors, enhances binding to F-actin.

### Non-muscle Tpm isoforms can form heterodimers *in vitro* and in cells

It has long been known that muscle Tpm isoforms function as a mixture of homodimers and heterodimers (81). Similarly, we have shown here that non-muscle Tpm isoforms purify from cells as both homodimers and heterodimers, and that their ability to heterodimerize is retained after boiling and cooling *in vitro*, with no apparent preference for one form of association vs. another. This is an important finding, since Tpm heterodimerization can further expand the range and functional diversity of Tpm-actin copolymers with segregated localization and function in cells. The prevalence of heterodimers in cells could be dictated by the time, location and amount of expression of one isoform vs. another.

In summary, we have developed a novel method for the expression of fully native proteins in human cells, we have demonstrated its applicability to two types of proteins of general interest, we have used this method to make new discoveries about non-muscle Tpm that had so far resisted investigation, and we have provided the scientific community access to our newly-developed vectors to facilitate the adoption of this method.

## Materials and methods

### Human cell expression vectors

#### Vector pJCX4

The QuickChange site-directed mutagenesis kit (Agilent Technologies, Santa Clara, CA) was used to remove two SapI restriction sites from vector pEGFP-C1 (Takara Bio USA, Mountain View, CA). A Woodchuck hepatitis virus (WHP) posttranscriptional regulatory element (WPRE) was added between the XbaI and MfeI sites of the multi-cloning site (MCS). The MCS-intein-chitin binding domain of vector pTXB1 (New England Biolabs, Ipswitch, MA) was then inserted between sites NheI and Kpn2I, and the MCS of vector pEGFP-C1 was removed by annealed oligos which were ligated between sites Kpn2I and XbaI. The vector is depicted in figure 1*A*.

#### Vector pJC8

The intein-CBD cassette was removed from vector pJCX4 and replaced with an internal ribosome entry site (IRES) element between sites BamHI and BspE1. Two new multicloning sites, one containing a V5- and the other a FLAG-affinity tag, were introduced on either side of the IRES using annealing oligos. The V5 and FLAG epitope tags provide clear signals by Western blot analysis and are used here to recognize the expressed Tpm isoforms. The MCSs and IRES region of the vector are depicted in figure 4*A*.

### Cloning of Tpm isoforms and α-synuclein

The following cDNAs were used in cloning (NCBI codes): Tpm1.6 (NM_001018004), Tpm1.7 (NM_001018006), Tpm2.1 (NM_213674), Tpm3.1 (NM_153649), Tpm3.2 (NM_001043351), Tpm4.2 (NM_003290), α-synuclein (NM_000345.4). The cDNAs were cloned into vector pJCX4 by PCR-amplification with primers that included the NotI (5’) and SapI (3’) restriction enzyme sites (*Figure 1, Figure S1*). The genes encoding isoforms Tpm1.6 and Tpm3.1 with added N-terminal Met-Ala-Ser in vector pBAT4 were a gift from Dr. Pekka Lappalainen. The two isoforms were also cloned without N-terminal extensions into vector pBAT4 by PCR-amplification with primers that included the NcoI (5’) and either BamHI or XhoI (3’) restriction enzyme sites. Four of the Tpm isoforms (Tpm3.1, Tpm3.2, Tpm3.5, Tpm4.2) were separately cloned as pairs into the IRES-containing vector pJC8 (see above) using PCR primers which included XhoI and BspEI (MCS1) or NotI and NheI (MCS2) on their overhangs. For this, the cDNA encoding isoform Tpm3.5 was generated starting from Tpm3.1 (amino acids 1-222) and the nucleotides encoding for the C-terminal 26 amino acids that differ among these two isoforms were added by overlap extension PCR. All the primers used in this study are listed in Table S2.

### Protein expression in Expi293F cells and intein-based purification

Expression was carried out with minor changes to the manufacture’s protocol for the Expi293 Expression System Kit (Thermo Fisher Scientific, Waltham, MA). Expi293F cells were grown in Expi293 media to a density of 2.5 x 10^6^ cells/mL. Cultures were incubated on an orbital shaker (125 rpm) at 37°C using an Isotemp incubator (Thermo Fisher Scientific, Waltham, MA) with a humidified atmosphere of 8% CO_2_. Pure plasmid DNA (1 µg per mL of Expi293F culture) and polyethylenimine (PEI, 3 µL per mL of Expi293F culture) were added to separate 5 mL aliquots of 37°C Opti-MEM (Thermo Fisher Scientific), mixed after 5 min, and incubated for 30 min at 25°C and then added to the Expi293F cell culture. Cultures were grown for 48-72 h, optimized for each protein. Cells were harvested by centrifugation (4,000 x g) at 4°C for 10 min.

For purification, cell pellets were resuspended in 20-35 mL of lysis buffer (20 mM HEPES pH 7.5, 500 mM NaCl, 1 mM DTT). Cells were lysed using a Dounce glass piston homogenizer (Kontes Glass, Vineland, NJ) and the supernatants were clarified by centrifugation at 48,000 x g at 4°C for 20 min. The clarified supernatants were loaded onto 15 mL chitin resin (packed into a column) pre-equilibrated with lysis buffer and washed extensively. Self-cleavage of the intein was activated with the addition of 50 mM β-mercaptoethanol at 25°C for 24 h. The expressed proteins were then eluted with lysis buffer, dialyzed overnight into A-buffer (20 mM HEPES pH 7.5, 50 mM NaCl) and further purified on a Source Q column (MilliporeSigma, St. Louis, MO) using a 50-300 mM NaCl gradient. Peak fractions were dialyzed into storage buffer (20 mM HEPES pH 7.5, 50 mM NaCl), flash frozen in liquid nitrogen and stored at -80°C.

### *E. coli* expression and purification of Tpm isoforms

Tpm isoforms Tpm1.6 and Tpm3.1 (with and without Met-Ala-Ser at the N-terminus) were expressed in ArcticExpress(DE3) RIL cells (Agilent Technologies), grown in Terrific Broth (TB) medium for 6 h at 37°C to an optical density of ∼1.5-2 at 600 nm (OD_600_), followed by 24 h at 10°C in with 0.4 mM isopropyl-β-D-thiogalactoside (IPTG). Cells were harvested by centrifugation (4,000 x g) and resuspended in 20 mM HEPES (pH 7.5), 100 mM NaCl, 1 mM EDTA, and 1 mM phenylmethanesulfonyl fluoride (PMSF). Cells were boiled in a water bath for 10 min, cooled on ice for 10 min, and clarified by centrifugation at 48,000 x g for 20 min at 4°C. Sodium acetate (1 M, pH 4.5) was added to the clarified supernatant containing the Tpm isoforms (that are heat-stable) until the solution reached the isoelectric point for precipitation of each isoform (4.5-4.7). The solutions were then centrifuged at 48,000 x g to pellet the Tpm isoforms. Pellets were dissolved in A-buffer (20 mM HEPES pH 7.5, 100 mM NaCl) and dialyzed in the same buffer overnight. Proteins were loaded on Source Q ion exchange (MilliporeSigma) and eluted with a 100-500 mM NaCl gradient. Peak fractions containing pure proteins were dialyzed in storage buffer (20 mM HEPES pH 7.5, 50 mM NaCl), flash frozen in liquid nitrogen and stored at -80°C.

### Preparation of samples for proteomics analysis

Gel bands of the purified proteins were cut and destained with 100 mM ammonium bicarbonate/acetonitrile (50:50). The bands were reduced in 10 mM dithiothreitol and 100 mM ammonium bicarbonate for 60 min at 52°C. The bands were then alkylated with 50 mM iodoacetamide and 100 mM ammonium bicarbonate at 25°C for 1 h in the dark. The proteins were digested with enzymes (trypsin, chymotrypsin, or Glu-C) at 37°C for 12 h. Supernatants were removed and transferred to fresh tubes. Additional peptides were extracted from the gel by adding 50% acetonitrile and 1% TFA and shaking for 10 min. The supernatants were combined and dried. The dried samples were reconstituted in 0.1% formic acid for mass spectrometry analysis.

### Proteomics analysis by nano-LC-MS/MS

Peptides were analyzed on a Q-Exactive Orbitrap Mass Spectrometer attached to an Easy-nLC (Thermo Fisher Scientific) at 400 nL/min. Peptides were eluted with a 25 min gradient of 5-32% ACN followed by 5 min 90% ACN and 0.1% formic acid. The data-dependent acquisition mode with a dynamic exclusion of 45 sec was used for analysis. A full MS scan range of 350-1200 m/z was collected, with a resolution of 70 K, maximum injection time of 50 ms, and automatic gain control (AGC) of 1×10^6^. A series of MS2 scans were then acquired for the top 12 most abundant ions of the MS1 scans. Ions were filtered with charge 2–4. An isolation window of 2 m/z was used with the quadruple isolation mode. Ions were fragmented using higher-energy collisional dissociation (HCD), with a collision energy of 27%. Orbitrap detection was used with a scan range of 140-2000 m/z, resolution of 30 K, maximum injection time of 54 ms and AGC of 50,000. All the samples were analyzed using a multiplexed parallel reaction monitoring (PRM) method based on a scheduled inclusion list containing the target precursor ions. Full MS scans were acquired on the Orbitrap from 350–1200 m/z at a resolution of 60,000, using an AGC of 50,000. The minimum threshold was set to 100,000 ion counts. Precursor ions were fragmented with the quadrupole using an isolation width of 2 m/z units, a maximum injection time of 50 ms and AGC of 10,000.

### Proteomics MS Data Analysis Including Peptide Identification and Quantification

The mass spectrometry raw spectra were processed using Proteome Discoverer version 2.3 (Thermo Fisher Scientific). The spectra were searched against a database of target protein sequences using default setting: precursor mass tolerance of 10 ppm, fragment mass tolerance of 0.02 Da, enzymes specific cleavage sites, and up to two missed cleavages. Cysteine carbamidomethylation was set as a fixed modification, while methionine oxidation and N-terminal or lysine acetylation were set as variable modifications. The search results were filtered using the target-decoy approach with the false discovery rate (FRD) cutoff of < 1% at the peptide and protein levels. To identify PTMs, the raw data were re-processed with the program MetaMorepheus version 0.0.316 optimized for PTM analysis (94). The calibrate, G-PTM-D, and search tasks of the program were used with default parameters, including default fixed and variable modifications, precursor mass tolerance of 5 ppm, enzyme specific cleavage, up to two missed cleavages, and an FRD cutoff of < 1% at the peptide and protein levels.

### Cosedimentation assays

Actin filaments at a fixed concentration of 7 µM were incubated for 30 min at 25°C with increasing Tpm concentrations in cosedimentation buffer (20 mM HEPES pH 7.5, 200 mM NaCl, 5 mM MgCl_2_) and a total volume of 100 µL. The reactions were centrifuged for 30 min at 80,000 rpm (280,000 rcf) to pellet F-actin and any bound Tpm (*Figure 3A, Figure S3*). For each Tpm protein, the cosedimentation experiments were repeated three or four times (*Figure S3*). After centrifugation, 100 µL of the supernatants were mixed with 25 µL of 4x Laemmli SDS-PAGE loading buffer (S fraction). Pellets were gently washed with 100 µL of cosedimentation buffer, to remove any remaining soluble protein, and resuspended in 100 µL of cosedimentation buffer and mixed with 25 µL of 4x Laemmli SDS-PAGE loading buffer (P fraction). Identical volumes (7 µL) of the P and S fractions were analyzed on SDS-PAGE gels, Coomassie-stained, and the bands were densitometrically quantified with the program Image Lab (BIO-RAD). In the absence of actin filaments, all the Tpm isoforms were found in the S fraction, whereas most of the actin was found in the P fraction with or without Tpm (*Figure 3A, Figure S3*). In the presence of actin filaments, Tpm cosedimented with actin to different degrees. The free Tpm concentration (plotted on the x-axis) was determined by dividing the intensity of the Tpm band in the S fraction by the sum of the intensities of the Tpm bands in the S and P fractions and multiplied by the total Tpm concentration. The fraction of Tpm bound in the pellet was determined by dividing the intensities of the Tpm and actin bands in the pellet fraction. According to a commonly used convention (95,96), we plotted on the y-axis the fraction of maximal Tpm binding to F-actin, determined by normalizing the ratio of the intensities of the Tpm and actin bands in the pellet fraction to the plateau value for each Tpm sample (i.e. the maximum fraction of Tpm/actin in the pellet fraction). This allows to compare side-by-side Tpm isoforms with different F-actin-binding affinities by plotting the fraction of maximal binding vs. the free Tpm concentration and fitting the data to a Hill equation for cooperative binding. We used the program Prism 9.0.0 (GraphPad) for fitting with a “specific binding with Hill slope” model, which yielded the apparent dissociation constant (*K*_*app*_) and Hill coefficient for each interaction (*Figure 3E*).

### Detection of non-muscle Tpm heterodimers

Expi293F cell cultures (25 mL) expressing pairs of Tpm isoforms using the IRES-containing vector pJC8 (see above) were resuspended in FLAG lysis buffer (phosphate buffered saline, 1 mM EDTA, 1 mM PMFS, and 0.5% Triton X-100) and lysed by three cycles of freeze-thaw in liquid nitrogen. Lysates were clarified at 4°C by centrifugation at 48,000 x g for 20 min, and incubated with 100 μL of anti-DYKDDDDK (FLAG) resin (Thermo Fisher Scientific) for 2 h at 4°C. The resin was washed five times with 200 μL lysis buffer. FLAG-tagged proteins were released from the resin by incubation with buffer supplemented with 50 mM FLAG peptide (Genscript, Piscataway, NJ) for 30 min.

For boiling and refolding experiments, the proteins purified from the FLAG resin were heated for 10 min at 90°C and then allowed to cool to 25°C over 30 min. The proteins were incubated for 2 h at 4°C with 100 µL of V5 resin, washed five times with 200 μL lysis buffer, and directly used for analysis. Equal volumes (100 μL) of lysate, flowthrough, 1^st^ wash, last wash, FLAG elution, and V5 resin-bound proteins were mixed with 25 μL of 4x Laemmli SDS-PAGE loading buffer and analyzed by Western blotting using primary FLAG (Santa Cruz Biotechnology, Dallas, TX) and V5 antibodies (Thermo Fisher Scientific). Secondary ECL-linked anti-mouse and anti-rabbit antibodies (Cytiva Life Sciences) were applied and the blots were imaged using a Pierce ECL Plus Western Blotting Substrate (Thermo Fisher Scientific) on a G:BOX Chemi XR5 imager (Syngene, Frederick, MD).

## Acknowledgements

This work was supported by National Institutes of Health Grants R01 GM073791 and R01 MH087950 to RD and T32 AR053461 to PJC. We thank Pekka Lappalainen for the gift of vectors for the bacterial expression of tropomyosin isoforms. We thank the Quantitative Proteomics Resource Core at the Perelman School of Medicine (University of Pennsylvania), Hyoungjoo Lee, Joseph Cesare, and Michael Gilbert for help with the proteomics data acquisition and analysis. We thank Michael Shortreed for help with the use of the program MetaMorpheus.

## Author contributions

Peter J. Carman: conceptualization, resources, data analysis, validation, investigation, visualization, methodology, project administration, writing and review/editing

Kyle R. Barrie: methodology, review/editing

Roberto Dominguez: conceptualization, supervision, project administration, funding acquisition, writing and review/editing

---

**Table S2.**
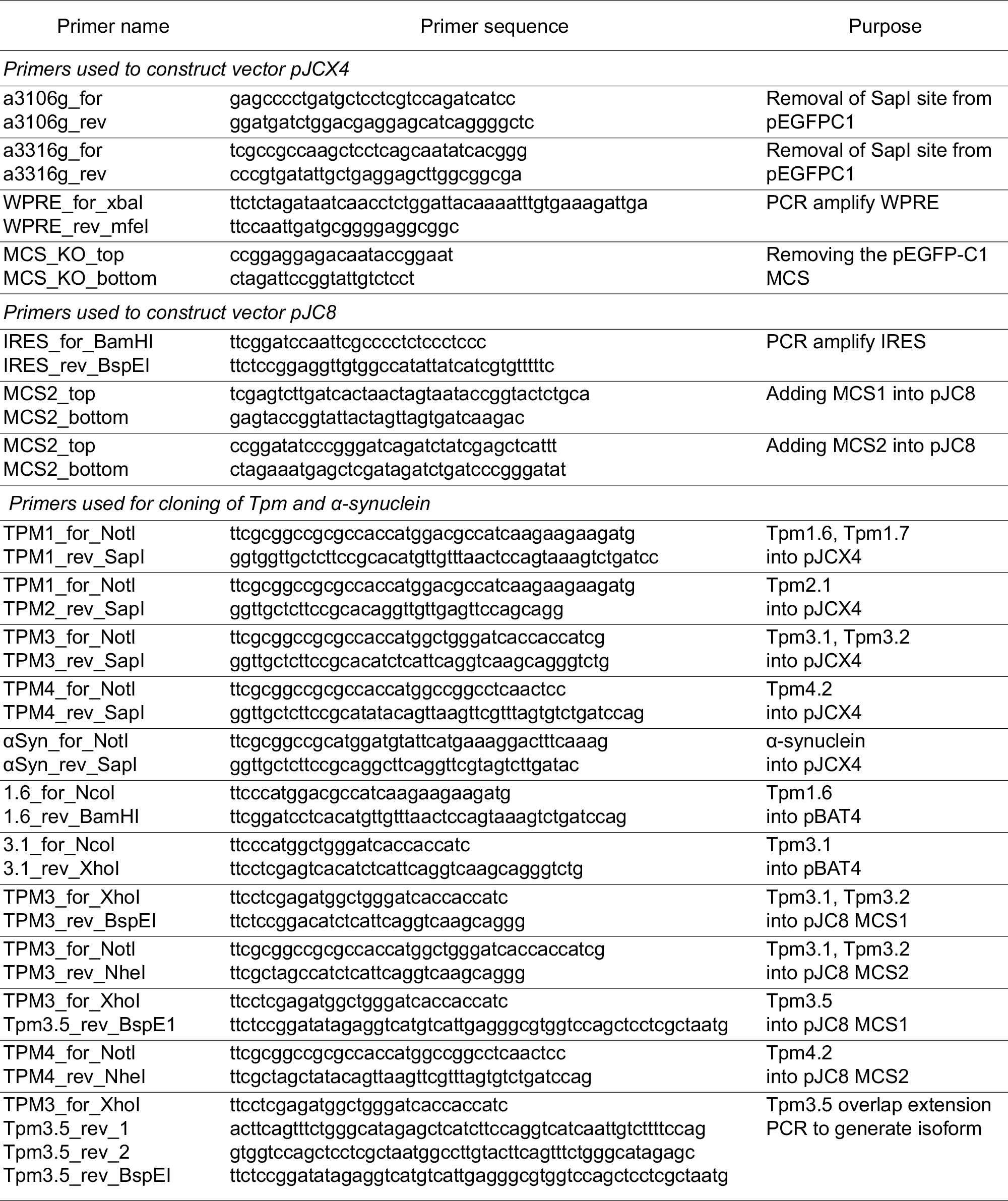
Primers used in this study

## Notes

### Competing Interest Statement

The authors have declared no competing interest.

### Summary of Updates

Figures on pages 4 and 5 were obscured by the header. They are resized here so they are no longer obscured.

